# Extending the gap and loading RecA: the presynaptic phase plays a pivotal role in modulating lesion tolerance pathways

**DOI:** 10.1101/2021.07.29.454318

**Authors:** Luisa Laureti, Lara Lee, Gaëlle Philippin, Michel Kahi, Vincent Pagès

**Affiliations:** Cancer Research Center of Marseille: Team DNA Damage and Genome Instability | CNRS, Aix Marseille Univ, Inserm, Institut Paoli-Calmettes, Marseille, France

## Abstract

During replication, the presence of unrepaired lesions results in the formation of single stranded DNA (ssDNA) gaps that need to be repaired to preserve genome integrity and cell survival. All organisms have evolved two major lesion tolerance pathways to continue replication: Translesion Synthesis (TLS), potentially mutagenic, and Homology Directed Gap Repair (HDGR), that relies on homologous recombination. In *Escherichia coli*, the RecF pathway repairs such ssDNA gaps by processing them to produce a recombinogenic RecA nucleofilament during the presynaptic phase. In this study, we show that the presynaptic phase is crucial for modulating lesion tolerance pathways. Indeed, impairing either the extension of the ssDNA gap (mediated by the nuclease RecJ and the helicase RecQ) or the loading of RecA (mediated by the RecFOR complex) leads to a decrease in HDGR. We suggest a model where defects in the presynaptic phase delay the formation of the D-loop and increase the time window allowed for TLS. We indeed observe an increase in TLS independent of SOS induction. In addition, we revealed an unexpected synergistic interaction between *recF* and *recJ* genes, that results in a *recA* deficient-like phenotype in which HDGR is almost completely abolished.

## INTRODUCTION

The genome of all living organisms is constantly damaged and some lesions can impair replication, potentially leading to mutagenesis and genome instability. In the presence of an unrepaired replication-blocking lesion, a single-stranded DNA (ssDNA) gap is formed, due to the initiation of the next Okazaki fragment (on the lagging strand) or to a repriming event (on the leading strand) (1–3). In order to complete replication and preserve cell survival, the ssDNA gap is filled in by one of the two lesion tolerance pathways, identified both in prokaryotes and eukaryotes: 1) Translesion Synthesis (TLS), which employs specialized DNA polymerases able to replicate the damaged DNA, with the potential to introduce mutations (4–6); 2) Damage Avoidance (DA) pathways, which use the information of the sister chromatid to bypass the lesion in a non-mutagenic way through homologous recombination mechanisms (7). In *Escherichia coli*, we have shown that in non-stressed conditions Homology Directed Gap Repair (HDGR), a DA mechanism that relies on the recombinase RecA, is the major lesion tolerance pathway employed by cells while TLS pathway is a minor one (8, 9). We have also identified another lesion tolerance strategy, named Damaged Chromatid Loss (DCL) that promotes cell proliferation at the expense of losing the damaged chromatid; in particular when HDGR mechanism is impaired, cells can survive by replicating only the undamaged chromatid (8).

In *E. coli* there are two well-known RecA dependent recombinational pathways: the RecBCD pathway, involved in the repair of double strand breaks, and the RecF pathway (also known as the RecFOR pathway), involved in the repair of ssDNA gap and required for an efficient SOS induction (reviewed in (10–12). Originally, the RecF pathway has received less attention because most of the studies on recombination focused on processes initiated by a double strand break, such as conjugation or transduction. However, in the last decades, it became clear that this pathway, other than be a “back-up” recombinational pathway in the absence of RecBCD, plays an important role during replication of a damaged DNA. Recently, it was shown that during non-stressed conditions, homologous recombination (RecA and RecF dependent) is mainly required to repair ssDNA gaps during replication other than double strand breaks (13).

The RecF pathway is evolutionary conserved in all bacteria (14) and functional orthologs are also found in Eukaria. This pathway encompasses several proteins and is divided into three distinct phases (11): i) the presynaptic phase in which the damaged ssDNA gap is processed by different proteins (*i*.*e*. SSB, RecF, RecO, RecR, RecJ, RecQ) to promote the formation of an active RecA nucleofilament; ii) the synaptic phase in which homology pairing and strand exchange by the RecA nucleofilament occur to produce a D-loop and iii) the postsynaptic phase involving the processing of the recombinational intermediates by the resolvase RuvABC or by the helicase RecG. Some of these proteins have already been well characterized *in vitro* (15–18), but their role *in vivo* and their impact on lesion tolerance pathways is not yet fully elucidated. The presynaptic phase is of particular interest since the ssDNA gap is the common substrate for both HDGR and TLS, hence how the ssDNA gap is processed to produce a RecA nucleofilament might influence the choice of lesion tolerance pathways.

Several genetic analyses indicate that the *recF, recO* and *recR* genes belong to the same epistasis group as the double mutants behave like the single mutants, at least for the UV sensitivity and the delay in SOS induction (19–23). According to *in vivo* and *in vitro* studies, it was proposed that two complexes, RecOR and RecFR, are formed whose function at a ssDNA gap is to disassemble the filament of single strand binding (SSB) proteins in order to load the recombinase RecA and promote homologous recombination (22, 23). *In vitro*, high amounts of the RecOR complex are sufficient to displace SSB proteins and load RecA, promoting strand exchange in the absence of RecF (24, 25). On the other hand, RecF was shown to bind to gapped substrates, preferentially on the 5’ double strand-single strand (ds-ss) DNA junction (23, 26) and often colocalize with the replisome (27). The UV sensitivity phenotype of the *recFOR* deficient strains is presumably due to a defect in RecA loading that leads to recombinational repair deficiency. Accordingly, a RecA mutant (*i*.*e*., the *recA730* allele) able to load itself on a ssDNA partially restores UV resistance of the *recFOR* deleted strains (28, 29).

The product of the *recJ* gene has also been associated with the RecF pathway (30); however, compared to the *recFOR* deficient strains, no strong UV sensitivity phenotype was observed when the *recJ* gene was deleted (19). Later, Courcelle and Hanawalt (31, 32) proposed that the products of *recJ* and *recQ* genes were involved in the repair of ssDNA gaps by processing the nascent lagging DNA strand during replication blockage. The helicase RecQ (3’->5’) would help the exonuclease RecJ (5’->3’) to enlarge the ssDNA at the 5’ end of the gap to lengthen the substrate for RecA filament formation. *In vitro* data showing that SSB protein is able to stimulate the activity of RecQ and of RecJ corroborate the hypothesis that both RecQ and RecJ are recruited at the ssDNA gap formed upon lesion encountering (18). Recently, we have shown that in the presence of an unrepaired lesion, RecJ is required to enlarge the ssDNA gap not only in the lagging strand but also in the leading strand (33), thus providing further evidence of the *in vivo* role of RecJ in the presynaptic phase.

In the present study, we have explored the role of the genes involved in the presynaptic phase of the RecF pathway, their genetic interaction and their effect on lesion tolerance pathways. We confirmed that, regarding lesion bypass, the *recF, recO* and *recR* genes belong to same epistasis group, likewise *recJ* and *recQ* genes showed to be epistatic. Defect in RecA loading (mediated by the RecFOR complexes) or in enlarging the ssDNA gap (mediated by RecQ and RecJ) results in HDGR impairment that is partially compensated by TLS independently of SOS induction. Interestingly, our data revealed a strong genetic interaction between *recF* and *recJ* (as well as *recF* and *recQ*), showing an additive effect of the deletion of both genes. Indeed, loss of these two genes results in a *recA* deficient-like phenotype in which HDGR is almost completely abolished, indicating that both the size of the ssDNA gap and the loading of RecA play an important role in the formation of an efficient D-loop structure, thus favoring the non-mutagenic HDGR mechanism over TLS pathway.

## MATERIAL AND METHODS

### Bacterial strains and growth conditions

All *E. coli* strains used in this work are listed in Supplementary Table 1. They are either derivative of strains FBG151 and FBG152 (9), that carry a plasmid that allows the expression of the *int–xis* genes after IPTG induction, or derivative of strains EC1 and EC2 (this study), in which the *int–xis* genes have been inserted into the chromosome of *E. coli*. Strains were grown on solid and in liquid Lysogeny Broth (LB) medium. Gene disruptions were achieved by the one-step PCR method (59) or by P1 transduction using the Keio collection (60). Following the site-specific recombination reaction, the lesion is located either in the lagging strand (FBG151 or EC1 derived strains) or in the leading strand (FBG152 or EC2 derived strains). Antibiotics were used at the following concentrations: ampicillin 50 or 100 μg/ml; tetracycline 10 μg/ml, kanamycin 100 μg/ml, chloramphenicol 30 μg/ml. When necessary, IPTG and X-Gal were added to the medium at 0.2mM and 80 μg/ml, respectively.

### Plasmids

pVP135 expresses the integrase and excisionase (*int–xis*) genes from phage lambda under the control of a *trc* promoter that has been weakened by mutations in the -35 and the -10 region. Transcription from Ptrc is regulated by the *lac* repressor, supplied by a copy of *lacI*^*q*^ on the plasmid. The vector has been modified as previously described (9).

pVP146 is derived from pACYC184 plasmid where the chloramphenicol resistance gene has been deleted. This vector, which carries only the tetracycline resistance gene, serves as an internal control for transformation efficiency.

pVP141-144 and pGP1, 2 and 9 are derived from pLDR9-attL-lacZ as described in (9). pLL1, pLL2c and pLL7 are derived from pVP141 and contain several genetic markers as previously described (8). All these plasmid vectors contain the following characteristics: the ampicillin resistance gene, the R6K replication origin that allows plasmid replication only if the recipient strain carries the *pir* gene, and the 5’ end of the *lacZ* gene in fusion with the *attL* site-specific recombination site of phage lambda. The P’3 site of *att*L has been mutated (AATCATTAT to AAT<u>TA</u>TTAT) to avoid the excision of the plasmid once integrated. These plasmids are produced in strain EC100D pir-116 (from Epicentre Biotechnologies, cat# EC6P0950H) in which the pir-116 allele supports higher copy number of R6K origin plasmids. Vectors carrying a single lesion for integration were constructed as previously described following the gap-duplex method (9). A 13-mer oligonucleotide, 5′-GCAAG<u>TT</u>AACACG-3′, containing no lesion or a TT(6-4) lesion (underlined) in the *Hinc*II site was inserted either into the gapped-duplex pLL1/2c leading to an out of frame *lacZ* gene (to measure HDGR) or into the gapped-duplex pGP1/2 leading to an in frame *lacZ* gene (to measure TLS0). A 15-mer oligonucleotide 5’-ATCACCGGC<u>G</u>CCACA-3’ containing or not a single G-AAF adduct (underlined) in the *Nar*I site was inserted into the gapped-duplex pLL1/7 (to measure HDGR) or into the gapped-duplexes pVP141-142 or pVP143-144 to score respectively for TLS0 Pol V-dependent and for TLS-2 Pol II-dependent. A 13-mer oligonucleotide, 5′-GAAGACCT<u>G</u>CAGG, containing no lesion or a dG-BaP(-) lesion (underlined) was inserted into the gapped-duplex pVP143/pGP9 leading to an in frame *lacZ* gene (to measure TLS).

### Monitoring HDGR and TLS events

The protocol for lesion integration assay is described in details in (39). Cells were diluted and plated before the first cell division using the automatic serial diluter and plater EasySpiral Dilute (Interscience) and were counted using the Scan 1200 automatic colony counter (Interscience). No differences were observed when we used FBG or EC derivative strains.

Following the integration of the pLL1/2c vector (TT6-4 lesion) or pLL1/7 vector (AAF lesion), sectored blue/white colonies represent HDGR events. Following integration of the pVP141/142, pVP143/144, pGP1/2, pVP143/pGP9 vectors (AAF, TT6-4 or BaP lesion, respectively), sectored blue/white colonies represent TLS events. The relative integration efficiencies of lesion-carrying vectors compared with their lesion-free homologues, normalized by the transformation efficiency of pVP146 plasmid in the same electroporation experiment, allow the overall rate of lesion tolerance to be measured. For each lesion we combine the results obtained with the different plasmids to measure HDRG and TLS events (for BaP lesion we only measure TLS events).

The data in every graph represent the average and standard deviation of at least three independent experiments of a lesion inserted in the leading (or in the lagging) orientation. Tolerance events (Y axis) represent the percentage of cells able to survive in presence of the integrated lesion compared to the lesion-free control.

## RESULTS

In the present work, we set out to investigate the presynaptic phase of the RecF pathway during the encounter of a replication fork with a DNA lesion. We used a previously described assay that allows to insert a single replication-blocking lesion into the chromosome of *E. coli* and monitor lesion tolerance mechanisms by a colorimetric assay based on the *lacZ* reporter gene (8, 9, 34). Briefly, a non-replicative plasmid containing a defined DNA lesion in the 5’-end of the *lacZ* gene is inserted into a precise locus of *E. coli* genome reconstituting the entire *lacZ* gene (Fig 1A). Plasmid integration is irreversible and is achieved by the site-specific recombination system of the bacteriophage lambda, integration events are selected using an antibiotic resistance marker. Using different types of lesion-containing plasmids, we can directly monitor either TLS (Fig 1B) or HDGR mechanisms (Fig 1C). For this study, we used two blocking lesions: the UV-induced thymine-thymine pyrimidine (6-4) pyrimidone photoproduct (TT6-4) and the chemical guanine adduct, the N-2-acetylaminofluorene (G-AAF). *E. coli* possesses three TLS polymerases, involved in the bypass of different types of DNA lesions: Pol II (encoded by *polB*) (35), Pol IV (encoded by *dinB*) (36) and Pol V (encoded by *umuDC*) (37). The expression level of these TLS polymerases is strongly increased upon SOS induction. UV lesions are exclusively bypassed by Pol V while the G-AAF lesion when positioned in a particular sequence context (*i*.*e*., the *Nar*I hotspot sequence) can be bypassed either by Pol V (named TLS0 events) or Pol II (named TLS-2 events, frameshift -2) (4, 38). The *E. coli* strains used in this study are deficient for the mismatch repair system (*mutS*) to prevent corrections of the genetic markers of the integration plasmids, as well as for nucleotide excision repair (*uvrA*), to avoid excision of the lesion and to focus on lesion tolerance events (see also (8, 9)). In our genetic assay, we can insert the lesion either on the lagging or in the leading orientation regarding the replication fork direction. Since we did not observe any significant difference between the two strand orientations, we will present in the manuscript the results obtained for a lesion inserted in the leading strand, while lagging strand results will be presented in Suppl Fig 1.

**Figure 1:**
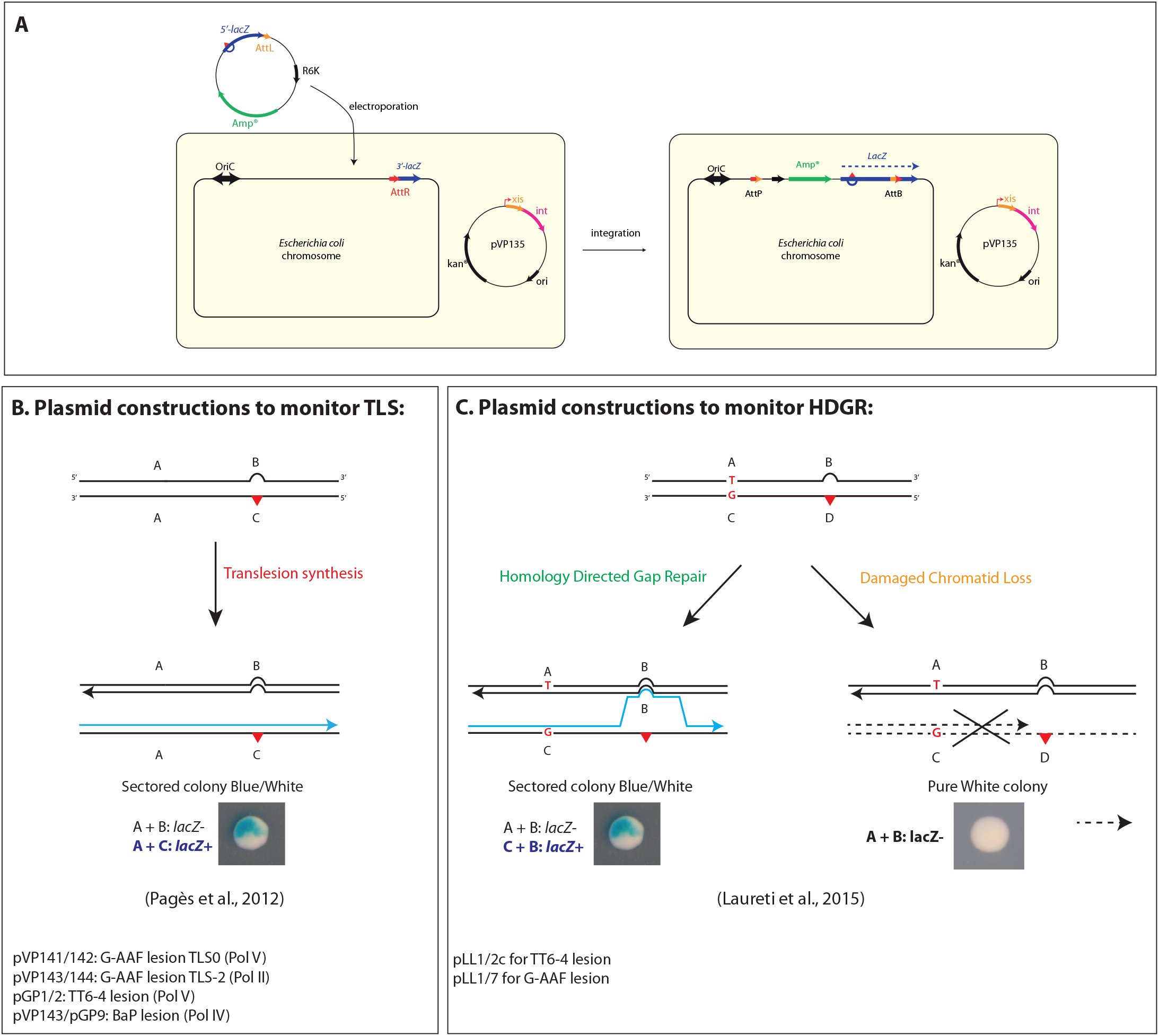
Lesion tolerance assay and lesion containing plasmids. **A)** The system is based on the phage lambda site-specific recombination and it requires a recipient *E. coli* strain with a single *att*R site fused to the 3’-end of the *lacZ* gene and a non-replicating plasmid construct containing a single DNA lesion (red triangle) in the 5’-end of the *lacZ* gene fused to the *att*L site. The recombination reaction between *att*L and *att*R is controlled by ectopic expression of phage lambda *int–xis* proteins (provided by pVP135), and leads to the integration of the lesion-containing vector into the chromosome. Integrants are selected on the basis of their resistance to ampicillin and the chromosomal integration restores an entire *lacZ* gene. **B)** Plasmid duplexes used to monitor TLS events (see also Pagès *et al*., 2012). The non-damaged strand (A+B markers) contains a short sequence heterology opposite the lesion that inactivates the *lacZ* gene and serves as a genetic marker for strand discrimination. Only the replication by TLS polymerases of the A+C markers will give rise to a functional *lacZ* gene and therefore to a sectored blue-white colony. For the G-AAF lesion, one construct measures TLS0 events specific of Pol V-dependent TLS events, while another one contains a +2 frameshit that allows to measure TLS-2 events specific of Pol II when the G-AAF lesion is inserted in the *NarI* hotspot sequence (see Pagès and Fuchs, 2002). **C)** Plasmid duplexes used to specifically monitor strand exchange mechanisms, *i*.*e*. HDGR events (see also Laureti *et al*., 2015). The system is a modified version of the plasmid constructs used to monitor TLS events. In these plasmids four genetic markers have been designed in order to distinguish the replication of the non-damaged strand (containing markers A and B) from the damaged strand (containing markers C and D). Using a combination of frameshift and stop codon, we inactivated *lacZ* gene on both the damaged (C-D) and undamaged (A-B) strands of the vector. Only a strand exchange mechanism by which replication has been initiated on the damaged strand (incorporation of marker C), and where a template switch occurred at the lesion site (leading to incorporation of marker B) will restore a functional *lacZ* gene (the combination of markers C and B contains neither a stop codon nor a frameshift); therefore leading to sectored blue-white colonies. When the damaged strand is loss, only the A and B marker will be replicating giving rise to a white colony.

### Loading of RecA: role of the mediators

In a previous study, we have shown that in the presence of a TT6-4 lesion the single mutants *recF* or *recO* but also the double mutant *recF recO* strongly reduced HDGR (Fig 2) (39). In the absence of its mediators (*i*.*e*. RecF, RecO or RecR), RecA loading can still occur, but randomly and with a slower kinetics (40, 41). Since HDGR relies on the recombinase RecA (8), this mechanism is necessarily affected. Similarly, induction of the SOS response is strongly affected by the delay in RecA nucleofilament formation (20, 21, 42). We measured a very low level of Pol V dependent TLS event (<0.5%) because there is essentially no Pol V expression in non-SOS induced cells (43). In addition, Pol V requires interaction with the tip of a RecA nucleofilament formed by RecFOR complex which less efficiently formed in the *recF* strain (44, 45).

**Figure 2.**
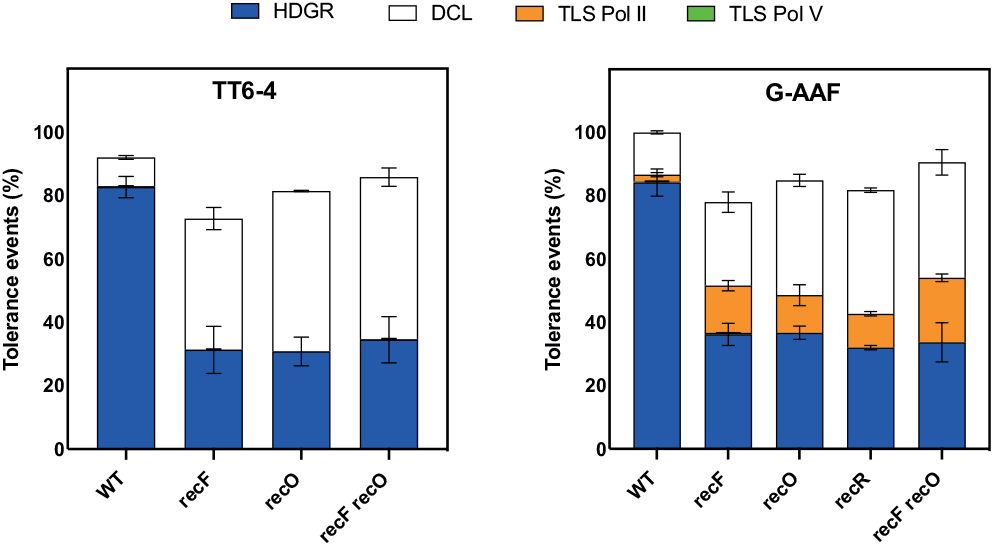
Deficiency in RecA loading promotes replication through TLS polymerases. The graphs represent the partition of lesion tolerance pathways (*i*.*e*. HDGR, TLS and DCL) for the single mutants *recF, recO* and *recR* or the double mutant *recF recO* in the presence of a TT6-4 lesion* (left panel) or a G-AAF lesion (right panel). All the strains are compared to our parental strain (WT in the graphs) which is *uvrA* and *mutS* deficient. *The data for the TT6-4 lesion have been previously published (39).

In the presence of a G-AAF lesion, deletion of either *recF, recO* or *recR* resulted again in a strong decrease in HDGR events, accompanied this time by a significant increase in Pol II dependent TLS events (Fig 2). The double mutant *recF recO* showed no further decrease in the level of HDGR compared to the single mutants, neither a significant increase in the level of Pol II dependent TLS (Fig 2). Altogether our genetic data indicate that the *recF, recO* and *recR* genes belong to the same epistasis group for lesion bypass and that the three genes play an important role in HDGR, which is expected due to their role in mediating RecA loading.

In a previous study, we have shown that affecting HDGR favors TLS: mutant alleles of RecA, that can form a nucleofilament but are impaired for homology search and strand exchange (*i*.*e*. are impaired in the D-loop formation), strongly favor lesion bypass by TLS polymerases (46). However, in this particular context, SOS response was constitutively activated thus the amount of TLS polymerases in the cells was significantly increased, which contributed to shifting the balance towards TLS pathway (46). Interestingly, in the *recF* (as well as *recO* or *recR*) deficient strain, despite the absence of SOS induction (SOS response is reduced and strongly delayed in these strains (20, 21)), the level of Pol II TLS at a G-AAF lesion increases to a level similar to the one observed after full SOS induction (either by deleting *lexA* or by UV-induction) (Suppl Fig 2). Our data point out that TLS can be regulated by factors other than the SOS response: defect in RecA loading is indeed sufficient to strongly increase the level of Pol II dependent TLS events, without activating the SOS response. To further confirm this, we monitored Pol IV activity on a benzo(a)pyrene (BaP): we show that the level of TLS in a *recF* deficient strain is again increased to the level observed upon full SOS induction by UV-irradiation of the cells (Suppl Fig 2).

### Effect of the ssDNA gap extension on lesion bypass

We have recently shown that the size of the ssDNA gap can modulate lesion tolerance pathways. Lack of the nuclease RecJ results in a shorter RecA-covered ssDNA gap, thus homology search and strand exchange are less efficient, impacting HDGR mechanism on one hand and favoring TLS pathway on the other hand (Fig 3 and (33)). While we have previously shown this for the G-AAF lesion, we confirmed that deletion of *recJ* affects HDGR mechanism also for the TT6-4 lesion (Fig 3).

**Figure 3.**
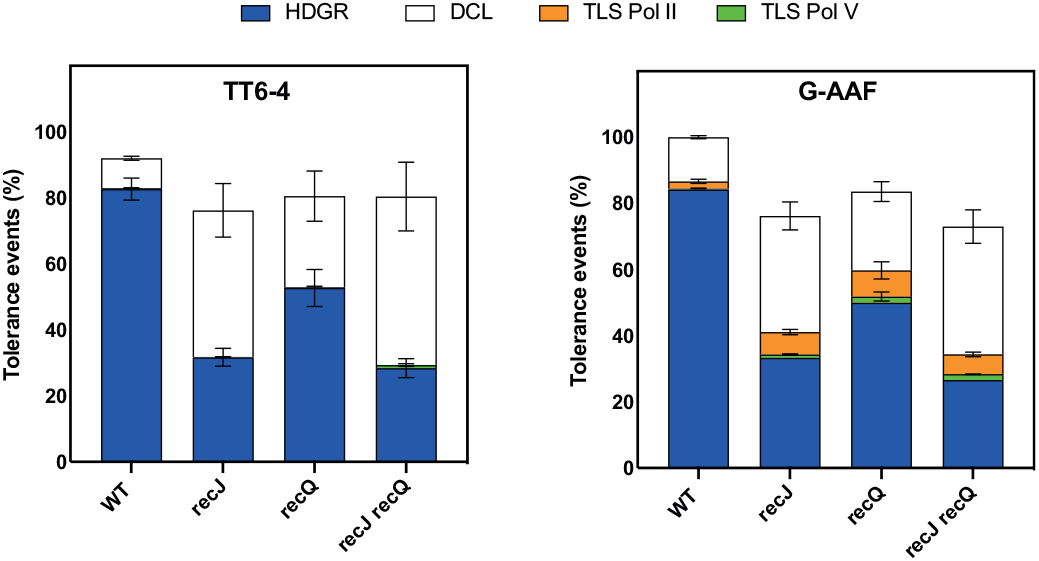
The size of the ssDNA gap affects the efficiency of HDGR mechanism favouring TLS. The graphs represent the partition of lesion tolerance pathways (*i*.*e*. HDGR, TLS and DCL) for the single mutants *recJ\** and *recQ* as well as for the double mutant *recJ recQ* in the presence of a TT6-4 lesion (left panel) or a G-AAF lesion (right panel). All the strains are compared to our parental strain (WT in the graphs) which is *uvrA* and *mutS* deficient. *\**The data for *recJ* in the presence of the G-AAF lesion have been previously published (33).

The helicase RecQ has been proposed to help RecJ in enlarging the ssDNA gap at the 5’ end (31). To confirm this role, we deleted the *recQ* gene and monitor lesion tolerance pathways in the presence of both G-AAF and TT6-4 lesions. We also observed a decrease in HDGR, but less important than when we deleted *recJ* or *recF* (Fig 3). This might indicate that even if the helicase activity of RecQ is required to provide the substrate for the nuclease activity of RecJ, it is not essential or that another helicase, yet to be identified, can partially compensate for the loss of RecQ, as also proposed by Courcelle *et al*. (32). In the presence of a G-AAF lesion, we measured an increase in Pol II dependent TLS similar to the *recJ* strain (Fig 3).

Deletion of both *recJ* and *recQ* in the presence of either a G-AAF or a TT6-4 lesion showed the same level of lesion tolerance pathways (for both HDGR and TLS) as the single mutant *recJ*. Hence, from our genetic analyses, *recJ* and *recQ* appeared to belong to the same epistasis group for lesion bypass, however RecJ plays a more critical role in HDGR mechanism.

It is interesting to note that in the *recJ, recQ* and *recJ recQ* mutants we observe a slight increase in Pol V TLS events (Fig 3 and Suppl Fig 3). In these strains, unlike in the *recF* deficient strain, RecA loading and nucleofilament formation occurs efficiently allowing the contact with Pol V that is required for its activation (44). This increase is only measured with the G-AAF lesion most likely because the low physiological level of Pol V is a limiting factor in the presence of the strong blocking lesion TT6-4. In line with our findings, Courcelle and coworkers observed in UV irradiated *E. coli* cells that Pol V was required for survival in the absence of *recJ* or *recQ* and that the increase of mutagenesis was Pol V dependent (32, 47).

Altogether, these results further confirm that impairing the extension of the ssDNA gap favors replication by TLS polymerases.

### Additive effect of *recF* and *recJ* genes

From our data, it appears that both RecA loading and the size of the ssDNA gap are important factors to modulate the partition and the efficiency of lesion tolerance pathways. To further investigate the presynaptic phase, we decided to analyze the genetic interaction between *recF* and *recJ*, by deleting both genes and monitoring lesion bypass in the presence of a G-AAF and TT6-4 lesion. Interestingly, for both lesions the double mutant showed a drastic drop in HDGR events (resulting in less than 10% of HDGR), compensated mainly by damaged chromatid loss (Fig 4). Similar results were obtained for the double mutant *recF recQ* (Fig 4). These data revealed that *recF* and *recJ* (or *recQ*) genes are not epistatic as it was expected since they are part of the same recombinational RecF pathway (30). On the contrary, deletion of both genes has an addictive effect on HDGR mechanism. No further effect was instead observed for TLS events: the additional loss in HDGR is compensated by an increase in DCL, not by a further increase in TLS. This is probably due to the fact that in the *recF* strain, the level of TLS already reaches that of a SOS induced strain (Suppl Fig 2) and cannot increase further. Likewise, no significant increase in Pol IV dependent TLS was observed when we deleted both *recF* and *recJ* genes in the presence of a BaP lesion (Suppl Fig 2).

**Figure 4.**
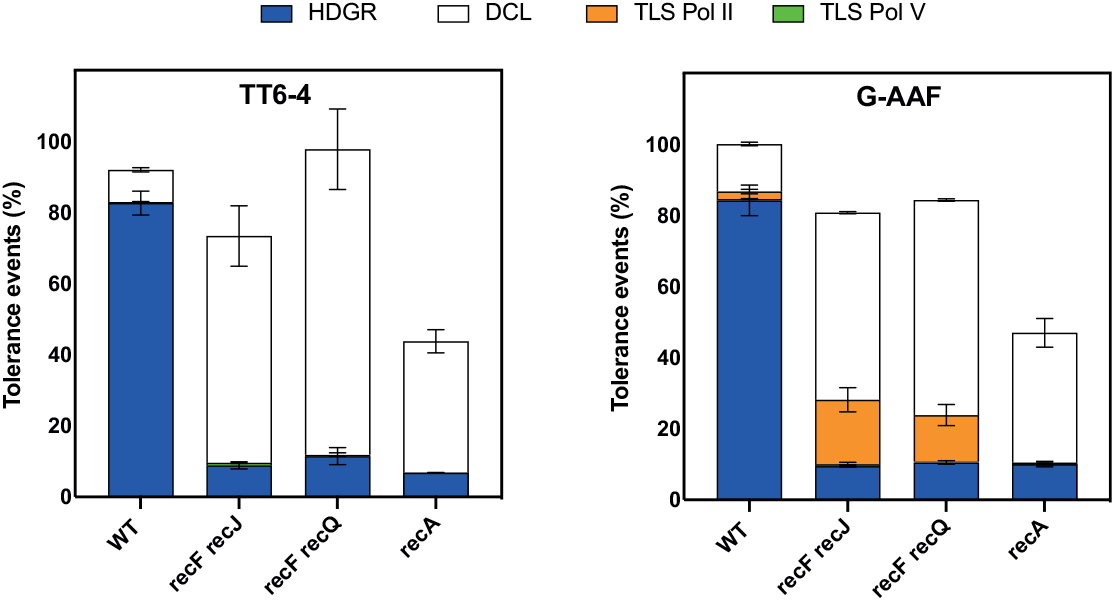
Additive effect of *recF* and *recJ* results in loss of HDGR mechanism. The graphs represent the partition of lesion tolerance pathways (*i*.*e*. HDGR, TLS and DCL) for the double mutant *recF recJ* and *recF recQ* in the presence of a TT6-4 lesion (left panel) or a G-AAF lesion (right panel). All the strains are compared to our parental strain (WT in the graphs) which is *uvrA* and *mutS* deficient. The data for the *recA* deficient strain have been already published (8).

A similar phenotype, *i*.*e*. a strong loss of HDGR, was previously observed in a *recA* deficient strain (see Fig 4 and also (8)). However, in the absence of RecA, we do not observe an increase in TLS. This might be explained by the fact that in the absence of RecF, RecA loading can still occur (but less efficiently) and this is sufficient to help stabilizing the ssDNA gap favoring replication by Pol II TLS. Similarly, a better stability of the ssDNA gap could explain why we did not observe a drastic decrease in cell survival for the double mutant *recF recJ* as in a *recA* deficient strain. To note, the 10% of sectored blue-white colonies still observed in the *recF recJ* (and *recF recQ)* strain as well as in the *recA* deficient strain result from a still uncharacterized RecA-independent mechanism, as previously proposed by the Rosenberg laboratory (13, 48).

Altogether these results indicate that both the loading of RecA and the size of the ssDNA gap are equally important for an efficient HDGR. Indeed, the simultaneous deficiency of these two processes condemns cells to survive mostly on damaged chromatid loss.

## DISCUSSION

In this study we investigated the role of the presynaptic phase of the RecF pathway on lesion bypass. The presynaptic phase is involved in the processing of the ssDNA gap formed upon encountering a lesion, and ends with the formation of a RecA nucleofilament. Once formed, the nucleofilament will engage in homology search and strand pairing, resulting in a D-loop structure that promotes HDGR and prevents TLS. The use of our single lesion assay allows to finely explore what occurs in the presence of a single replication blocking lesion and to better decipher lesion bypass mechanisms.

The presynaptic phase of the RecF pathway comprises several proteins that were expected to belong to the same epistasis group (30). While we confirmed this for the *recF-recO-recR* genes as well as for *recQ*-*recJ*, we unveiled an additive effect between *recF* and *recJ* (and *recF-recQ*), that results in a drastic loss of HDGR mechanism reminiscent of a *recA* deficient phenotype. We also showed that a defect in RecA loading and nucleofilament formation, as in a *recF* strain, or a defect in ssDNA gap extension, as in *recJ* strain, not only strongly affects HDGR mechanism but also promotes replication by Pol II or Pol IV TLS in the absence of SOS induction (*i*.*e*. without increasing the amount of TLS polymerases).

In non-stressed conditions, tolerance of highly blocking lesions relies almost exclusively on HDGR while TLS is a minor pathway (8). We have already shown that several factors contribute to regulate the balance between HDGR and TLS. In bacteria, TLS is positively regulated by the SOS response that increases the amount of the TLS polymerases in the cell, but TLS can also be modulated by HDGR efficiency: RecA mutants impaired in D-loop formation show a defect in HDGR and an increase in TLS (46). Similarly, the proximity of lesions on opposite strands generates overlapping ssDNA regions that prevent HDGR, leading again to an increase in TLS (33). In this work, we show that it is specifically the presynaptic phase of the RecF pathway that is critical for lesion tolerance pathway choice. Indeed, impairing the presynaptic phase leads to a delay in the formation of an efficient D-loop and therefore increases the time window allowed for TLS to occur, leading to a significant increase in TLS.

Based on these data, we propose a model that illustrates how the choice of lesion tolerance pathways is directed during the presynaptic phase of the RecF pathway (Fig 5). In the presence of an unrepaired lesion, the replicative polymerase temporarily stalls before a repriming event allows replication to continue, leaving a ssDNA gap. SSB protein will immediately cover the ssDNA gap to protect it and to orchestrate the recruitment of several proteins. SSB was indeed shown to directly interact with most of the proteins of the presynaptic phase as well as with the TLS polymerases (49, 50). The SSB-coated ssDNA gap can be regarded as a platform for protein recruitment for the processing of the ssDNA gap and for lesion bypass. The 5’ ds-ssDNA junction of the ssDNA gap appears to be a shared substrate for both RecQJ and RecFR complexes. Whether and which factors regulate the timing of RecQJ and RecFR recruitment is not known yet, however, we suggest that RecQ and RecJ are the first to be recruited at the 5’ junction of the nascent DNA. We based this hypothesis on the fact that RecQ and RecJ were shown to directly interact with SSB in the presence of a ssDNA gap and this interaction stimulates their catalytic activity to enlarge the ssDNA gap (18, 51–53). Supporting our hypothesis, Xia and coworkers (13) showed that RecJ and RecQ promote recombination and Holiday junction formation in vegetative *E. coli* cells, acting upstream of RecA during the repair of ssDNA gaps. Therefore, the extension of the gap must come before the loading of RecA. We have previously estimated the size of the ssDNA gaps to be in the range of 1.8-3.5 Kb (33) which correlates with the processivity established *in vitro* for RecJ (52). Once the complex RecQJ falls off, RecFR complex can have access to the 5’ ds-ssDNA junction and this also allows the recruitment of the RecOR complex to the ssDNA gap where RecO directly interacts with SSB (54, 55). Finally, RecA loading and filament formation can start. Since it occurs preferentially in the direction 5’->3’ (56), it will inhibit a new RecQ-RecJ recruitment. Once the RecA nucleofilament on the ssDNA gap is formed it will engage in homology search and strand invasion, forming the so-called D-loop structure required for the HDGR mechanism, that we showed to be the major lesion tolerance pathway employed by cells under non-stressed conditions (8, 9).

**Figure 5.**
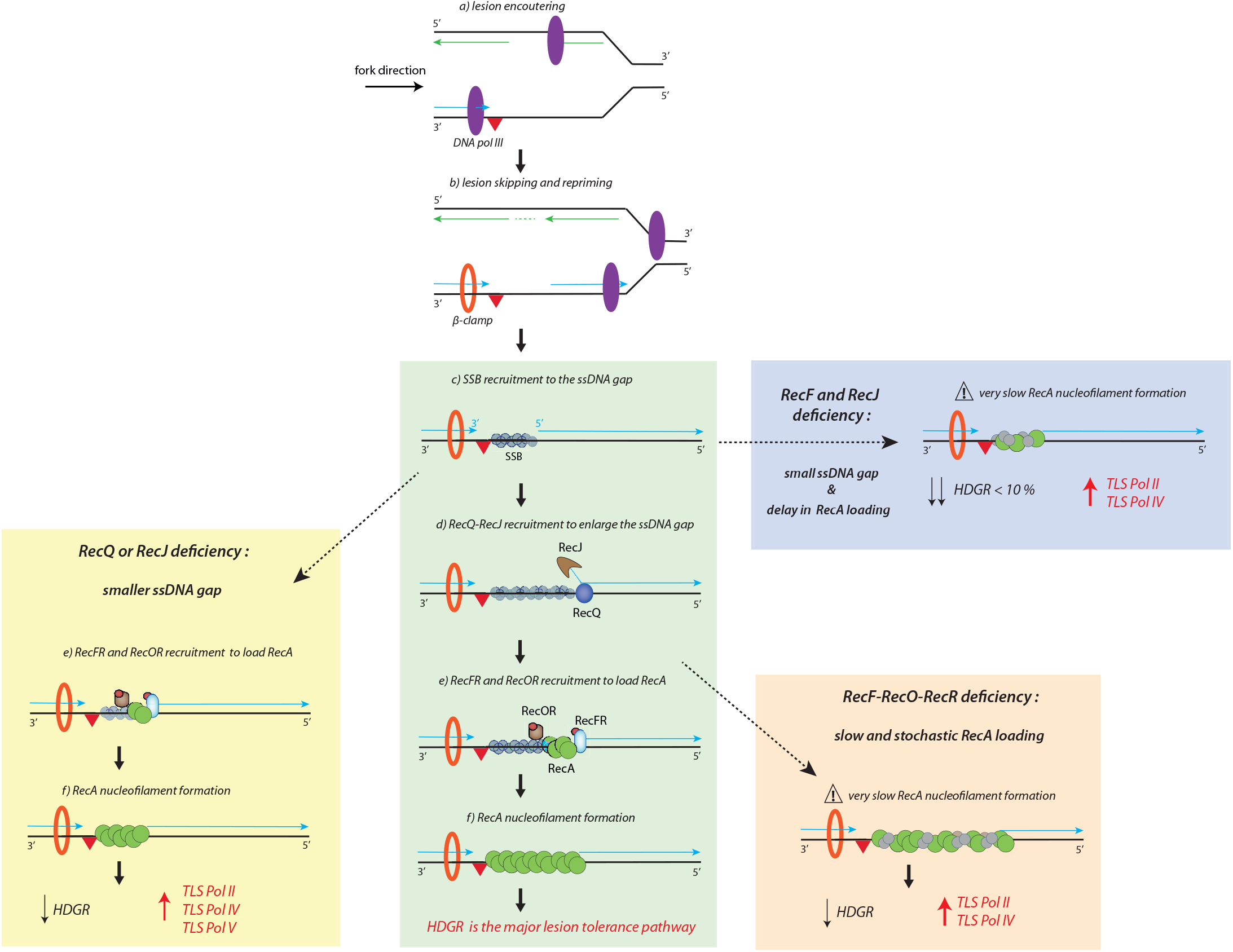
The presynaptic phase of the RecF pathway. Here is represented a scheme of the presynaptic phase during lesion bypass in the presence of a replication-blocking lesion (red triangle) in the leading strand. For simplicity after the repriming event (b), we have drawn only the molecular mechanisms occurring on the ssDNA gap. During replication, unrepaired lesions can temporarily block the replicative polymerase (a). The polymerase can skip the lesion and restart replication downstream, leaving a ssDNA gap (b). SSB protein is immediately recruited to the ssDNA gap (c) to protect it from undesired degradation, but also to orchestrate the recruitment of the proteins involved in the processing and the filling in of the ssDNA gap. The helicase RecQ and the nuclease RecJ are loaded at the 5’ ds-ssDNA junction and their concerted action enlarge the size of the gap (d). The processivity of RecJ-RecQ drops and they are replaced by the RecFR complex that is also recruited at the 5’ junction (e). The action of RecFR and of the RecOR complex at the ssDNA gap is to displace SSB and load the

Our data show that defect in the genes of the presynaptic phase affects HDGR mechanism favoring TLS pathway. Indeed, in the absence of either RecJ or RecQ, the size of ssDNA gap in not enlarged, thus impairing the efficiency of HDGR mechanism. Likewise, deletion of the RecA mediators (*recF, recO* or *recR*) causes a significant delay in RecA nucleofilament formation, as shown *in vitro* where nucleation time goes from 2-10 min in the presence of RecFOR to 10-60 min in its absence (40). In both cases, we observed an increase in TLS pathways. Absence of RecF, other than causing a delay in RecA filament formation, could allow RecJ to be recruited again and further extend the gap, a too long ssDNA gap can also affect HDGR mechanism. Indeed, in *recF* strain it was observed a RecJ dependent degradation of the nascent DNA that caused a delay in replication resumption (57, 58). Finally, the concomitant deficiency of RecF and RecJ results in a drastic decrease of HDGR mechanism, indicating that the slow formation of a small RecA-covered ssDNA strongly inhibits HDGR.

From our model, we can evince that the presynaptic phase, that includes the different steps to achieve the formation of the RecA nucleofilament, is crucial in modulating lesion tolerance pathways and in turn the outcome of lesion bypass (non-mutagenic versus mutagenic). The presynaptic phase can be considered as the “time window” where TLS can take place. Once the D-loop is formed, the recombinogenic RecA filament engages in HDGR process, thus closing the window that was allowed for TLS. Any delay in the presynaptic phase will extend this time window leading to an increase in TLS.

## ACKNOWLEDGEMENT

We thank Jean-Hugues Guervilly and Florencia Villafañez for critical reading of the manuscript.

