## Supplementary material for "Extending the gap and loading RecA: the presynaptic phase plays a pivotal role in modulating lesion tolerance pathways": suppl. fig

Supplementary data

**Supplementary Table 1: *E. coli* strains used in this study**

| Name | Short name | Genotype |
|---|---|---|
| EVP22/23 | Parental strain | FBG151/152 *uvrA*::frt *mutS*::frt |
| EVP146/147 | *recO* | FBG151/152 *uvrA*::frt *mutS*::frt *recO::*frt |
| EVP195/196 | *recQ* | FBG151/152 *uvrA*::frt *mutS*::frt *recQ* ::frt |
| EVP602/603 | *recO recF* | FBG151/152 *uvrA*::frt *mutS*::frt *recO::*frt *recF::*Kan |
| EVP347/348 | *recJ* | FBG151/152 *uvrA*::frt *mutS*::frt *recJ::frt* |
| EVP498/499 | *phrB recF* | FBG151/152 *uvrA*::frt *mutS*::frt *phrB* ::frt *recF* ::frt |
| EVP588/589 | *recJ recF* | FBG151/152 *uvrA*::frt *mutS*::frt *recJ*::frt *recF* ::frt |
| EVP615/616 | *recQ recF* | FBG151/152 *uvrA*::frt *mutS*::frt *recQ* ::frt *recF* ::frt |
| EVP963/964 | *recQ recJ* | FBG151/152 *uvrA*::frt *mutS*::frt *recQ* ::frt *recJ::frt* |
| EVP993/994 | *recR* | FBG151/152 *uvrA*::frt *mutS*::frt *recR*::frt |
| EVP989/990 | *phrB recF recO* | FBG151/152 *uvrA*::frt *mutS*::frt *phrB* ::frt *recF* ::frt *recO* ::frt |
| EC13/14 | Parental strain | EC1/EC2 *uvrA*::frt *mutS*::frt |
| EC39/40 | *recO* | EC1/EC2 *uvrA*::frt *recO::*frt *mutS*::frt |
| EC49/50 | *recO recF* | EC1/EC2 *uvrA*::frt *recO::*frt *mutS*::frt *recF* ::frt |
| EC51/52 | *recF* | EC1/EC2 *uvrA*::frt *recF::*frt *mutS*::frt |

**FBG151 =** MG1655 ΔlacIZ ΔattBλ::attRλ-3’lacZ-aadA (lesion inserted in the lagging orientation compared to replication)
**FBG152 =** MG1655 ΔlacIZ ΔattBλ::3’lacZ-attRλ-aadA (lesion inserted in the leading orientation compared to replication)
**EC1 =** MG1655 ΔlacI-lacZ::frt ΔattBλ::attRλ-3'lacZ-aad intergenic(aqpZ-ybjD)::high pTRC λ int-xis lacIq
**EC2 =** MG1655 ΔlacI-lacZ::frt ΔattBλ::3'lacZ-attRλ-aad intergenic(aqpZ-ybjD)::high pTRC λ int-xis lacIq

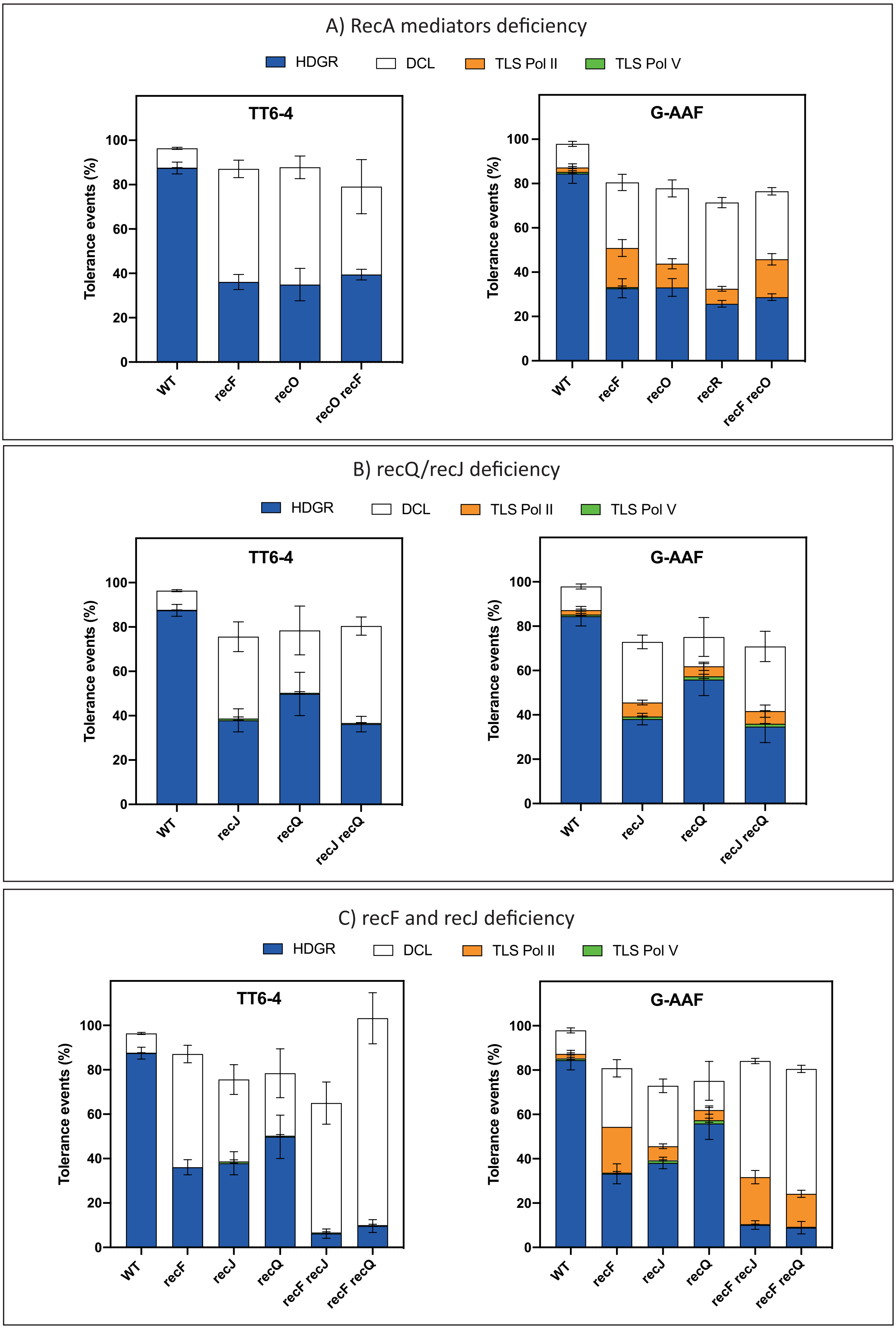


**Supplementary Figure 1:** The graphs represent the partition of lesion tolerance pathway (*i.e.* HDGR, TLS and DCL) in the presence of a TT6-4 lesion (on the left side) or a G-AAF lesion (on the right side) inserted in the lagging orientation. In panel **A)** are represented the data for the single mutants *recF, recO, recR* and the double mutant *recF recO*. In panel **B)** are represented the data for the single mutants *recJ, recQ,* and the double mutant *recJ recQ*. In panel **C)** are represented the data for the double mutants *recF recJ* and *recF recQ*. All the mutant strains are compared to our parental strain (WT in the graphs) which is *uvrA* and *mutS* deficient.

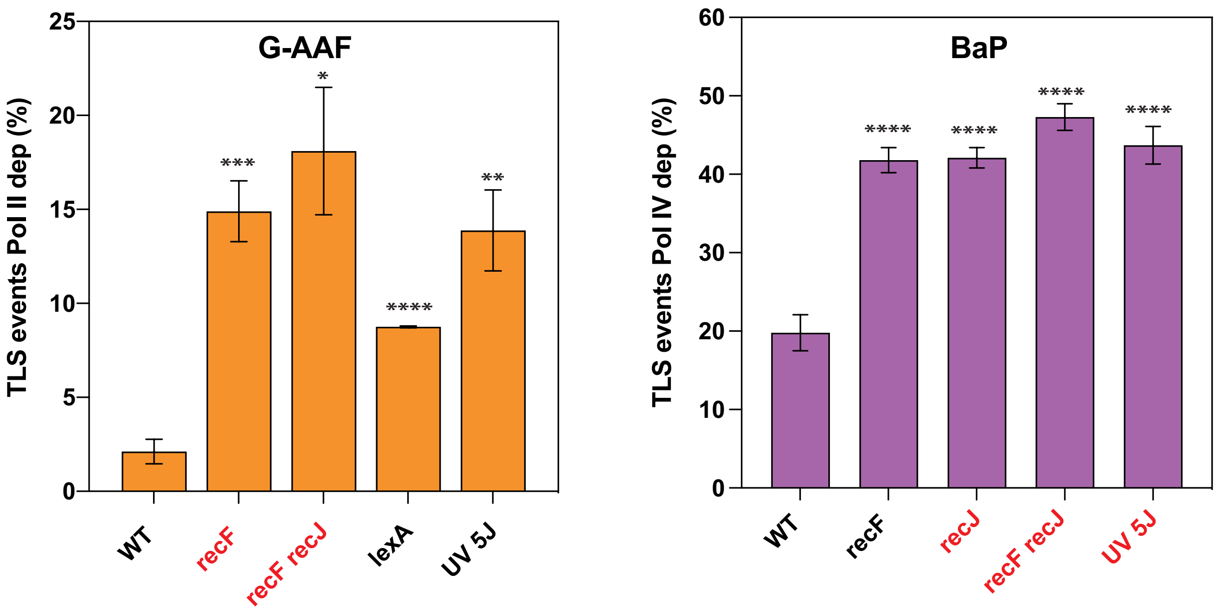


**Supplementary Figure 2:** The graphs represent the percentage of Pol II dependent TLS events in the presence of a G-AAF lesion (left panel) or the percentage of Pol IV dependent TLS events in the presence of a BaP lesion (right panel). All the mutant strains are compared to our parental strain (WT in the graphs) which is *uvrA* and *mutS* deficient. SOS induction is obtained either by deleting the *lexA* repressor gene or by UV induction (using 5J). The data in red have been obtained in this study, while the data in black have been previously published in (Naiman et al., 2014; Naiman et al., 2016). T-test was performed to compare values from the mutant strains to our parental strain (WT). *P < 0.05; **P < 0.005; ***P < 0005; ****P < 0.0001.

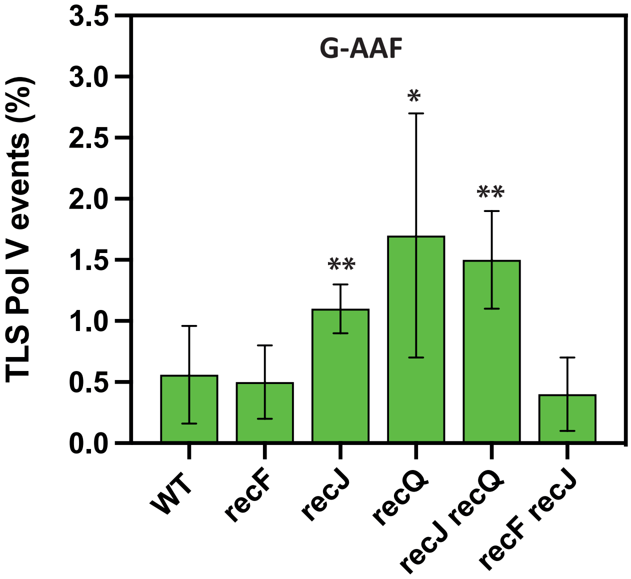


**Supplementary Figure 3:** The graph represents the percentage of Pol V dependent TLS events in the presence of a G-AAF lesion inserted either in the leading or in the lagging orientation (we pulled together the data since no difference was observed). There is a slight but statistically significant increase for the single mutants *recJ* and *recQ* as well for the double mutant *recJ recQ*. T-test was performed to compare values from the mutant strains to our parental strain (WT). *P < 0.05; **P < 0.005.
